# Cholecystokinin released somatodendritically from dopamine neurons broadly alters synaptic strength across the ventral tegmental area

**DOI:** 10.64898/2026.03.12.711406

**Authors:** Setareh Sianati, Yihe Ma, Julie A. Kauer

## Abstract

Neuropeptides are found in nearly every brain neuron, and can modulate behaviors by regulating neuronal excitability, synaptic transmission, and plasticity. In contrast to the canonical view of neuropeptide release from nerve terminals, we previously reported the somatodendritic release of cholecystokinin (CCK) from ventral tegmental area (VTA) dopamine (DA) neurons. Release of CCK occurs during modest depolarization of VTA DA cells, and by activating CCK2Rs, potentiates synaptic transmission from GABAergic afferents. Here, recording from dopamine neurons in acute midbrain slices from male and female mice, we examined how somatodendritic release of CCK regulates synaptic plasticity and the extent of its influence. Depolarization of a dopamine neuron induced long-term potentiation (LTP) at GABAergic synapses, and in parallel somatodendritic CCK release produced long-term depression (LTD) at glutamatergic synapses. CCK-induced LTP persisted when postsynaptic G protein signaling in dopamine neurons was blocked, suggesting that CCK likely acts at GABAergic presynaptic terminals. Activation of kappa opioid receptors prevented CCK-dependent LTP of GABAergic synapses, indicating interaction between these two neuromodulatory signaling pathways in VTA. Surprisingly, depolarization of one dopamine neuron potentiated synapses onto both the depolarized neuron and neighboring dopamine neurons located up to ∼100 µm away, indicating substantial spread of CCK signaling and synaptic modulation within the VTA region. Taken together, our findings demonstrate that somatodendritic CCK release bidirectionally coordinates synaptic strength across dopamine neurons, identifying a peptide-mediated feedback mechanism that shapes VTA circuit function.

**Significance Statement:** Dopamine neurons in the ventral tegmental area (VTA) play central roles in reward, motivation, stress responses, and feeding behavior. While fast synaptic inputs regulate dopamine neuron firing on fast timescales, less is known about how slower neuromodulatory signals shape these circuits. We show that somatodendritic release of the neuropeptide cholecystokinin from dopamine neurons coordinately alters both inhibitory and excitatory synaptic strength and influences neighboring neurons within the VTA. This peptide-mediated feedback mechanism operates over a broader spatial scale than classical synaptic transmission and is regulated by kappa opioid signaling. These findings reveal how local peptide release can reshape dopamine circuit function and may contribute to changes in reward processing and feeding behavior.

## Introduction

The VTA dopamine system plays a central role in reward processing, motivation, reinforcement learning and aversive behaviors (Wise, 2004; Lammel et al., 2012; Salamone and Correa, 2012; Margolis and Karkhanis, 2019). Dopamine neurons fire spontaneously, but their excitability is tightly regulated by the balance of inhibitory and excitatory synaptic inputs. Glutamatergic projections from the prefrontal cortex to the VTA increase burst firing of dopamine neurons (Gariano and Groves, 1988; Murase et al., 1993). In contrast, pharmacological and optogenetic studies have demonstrated that activation of different GABA afferents synapsing on dopamine neurons suppresses firing (Tan et al., 2012; Van Zessen et al., 2012; Polter and Kauer, 2014; Simmons et al., 2017; St Laurent et al., 2020).

While excitatory and inhibitory synaptic inputs provide rapid control of VTA dopamine neuron firing, these signals are embedded within a broader neuromodulatory framework that shapes excitability over longer timescales. Although classically studied during release from axon terminals, neurotransmitters can also be released from somatodendritic compartments. Somatodendritic release was first described for dopamine in rat substantia nigra (Geffen et al., 1976), and subsequent work demonstrated that somatodendritic dopamine release reduces dopamine cell excitability via D2 autoreceptors (D2R) (Beckstead et al., 2004; Ford et al., 2007). Somatodendritic release in multiple brain areas also occurs for several neuropeptides (Wagner et al., 1991; Simmons et al., 1995; Ludwig, 1998; Ludwig and Leng, 2006; Crosby et al., 2015; Tschumi et al., 2022; Caramia et al., 2023; Martinez Damonte et al., 2023; Hevesi et al., 2024), highlighting this form of signaling as physiologically significant. Signaling by somatodendritic release may either result in autocrine effects on the releasing neuron, or paracrine retrograde effects on synaptic afferents to the cell or region; both usually result from activation of G protein-coupled receptors. The best-studied examples of somatodendritic neuropeptide releasing neurons are the magnocellular vasopressin and oxytocin cells of the supraoptic nucleus, where both autocrine and paracrine effects have been documented (Ludwig and Leng, 2006). Notably, unlike point-to-point axon terminal communication with a postsynaptic site, somatodendritic neuropeptide release holds the potential for control of synaptic transmission across a broader region via volume transmission.

CCK is one of the most abundant neuropeptides in the brain and has been implicated in synaptic modulation across multiple brain regions (Crosby et al., 2015; Crosby et al., 2018; Chen et al. 2019), and broadly in mood regulation, learning, memory formation and the regulation of food intake (Derrien et al., 1994; Blevins et al., 2000; Voits et al., 2001; Chhatwal et al., 2009; Chen et al., 2019; Grove et al., 2025; Wang et al., 2025). We recently reported that somatodendritic release of CCK from VTA dopamine neurons selectively potentiates synaptic transmission from a subset of GABAergic afferents; a similar finding was also seen in the dorsomedial hypothalamic nucleus (Crosby et al., 2015). In VTA, modest depolarization (−40 mV for 6 min) or optogenetic activation of a single dopamine neuron is sufficient to trigger CCK release and potentiate GABAergic synaptic transmission, providing an autoinhibitory feedback mechanism on dopamine neurons (St Laurent et al., 2020; Martinez Damonte et al., 2023). In the substantia nigra, neurotensin similarly regulates dopamine neuron excitability, causing LTD of dopaminergic IPSCs (Tschumi et al., 2022).

Although somatodendritic and axonal release share several features, they are mechanistically distinct processes (Chen and Rice, 2001; Ludwig and Leng, 2006; Witkovsky et al., 2009; Kennedy and Ehlers, 2011; Rice and Patel, 2015; Hikima et al., 2021). Thus, defining the cellular mechanisms of somatodendritic CCK release is critical for understanding under what conditions peptide signaling regulates dopamine-neuron excitability. Here we investigate in more detail how somatodendritic release of CCK from VTA dopamine neurons regulates long-term synaptic plasticity at inhibitory and excitatory synapses, and how this signaling is influenced by a second neuromodulatory pathway, the kappa opioid receptor system. We also tested the spatial extent of CCK signaling within the VTA. We identify a peptide-mediated mechanism that bidirectionally coordinates synaptic strength within dopaminergic circuits.

## Methods

### Animals

All procedures were conducted in accordance with the National Institutes of Health guidelines for the care and use of laboratory animals and were approved by the Stanford University Administrative Panel on Laboratory Animal Care. Pitx3-GFP male and female mice were bred in-house and used in all experiments (Jackson Laboratory, B6.129P2-Pitx3*^tm1Mli^*/Mmjax, MMRRC ID: 41479) (Zhao et al., 2004).

### Preparation of brain slices

Acute brain slices were prepared as described previously (Martinez Damonte et al., 2023). Briefly, deeply anesthetized male and female Pitx3-GFP mice were perfused with an ice-cold, oxygenated choline-based solution containing (in mM): 110 choline chloride, 25 NaHCO_3_, 2.5 KCl, 1.25 NaH_2_PO_4_, 0.5 CaCl_2_, 7 MgCl_2_, 11.6 sodium ascorbate, 3.1 sodium pyruvate and 25 glucose. Following perfusion, horizontal slices (220 μm) were prepared using a Leica 1200 vibratome. Slices recovered at 34°C for 1 hour in oxygenated artificial cerebrospinal fluid (ACSF) containing in mM: 126 NaCl, 21.4 NaHCO_3_, 2.5 KCl, 1.2 NaH_2_PO_4_, 2.4 CaCl_2_, 1.2 MgSO_4_ and 11.1 glucose, saturated with 95% O_2_/5% CO_2_ (pH 7.4; 290-300 mOsm), and then were held submerged at room temperature until being transferred to the recording chamber.

### Electrophysiological recordings

Horizontal midbrain slices were continuously perfused (at 1.5-2 ml/min) with oxygenated ACSF at 29 – 30°C. Pitx3-GFP mice express GFP in Pitx3-expressing neurons, which overlap nearly 100% with tyrosine hydroxylase positive neurons of the VTA (Zhao et al., 2004). Whole-cell patch-clamp recordings were obtained from GFP-labeled neurons in the lateral VTA exhibiting a large hyperpolarization-activated current (I_h_>50 pA) in response to a voltage step from −40 mV to −120 mV. We used this criterion for consistency with our previous work; dopamine neurons exhibiting a large Ih current are most often located in the lateral VTA. Dopamine cells were voltage-clamped at −70 mV to minimize action potential firing during synaptic stimulation. To isolate GABAergic synaptic inputs, the AMPA receptor antagonist 6,7-dinitroquinoxaline-2,3-dione (DNQX; Tocris; 10μM) and the glycine receptor antagonist strychnine (Sigma-Aldrich; 1μM) were added to the ACSF. To isolate glutamatergic synaptic inputs, the GABA_A_ receptor antagonist bicuculline (Sigma-Aldrich; 10μM) and the glycine receptor antagonist strychnine (Sigma-Aldrich; 1μM) were added. Input resistance and series resistance were monitored throughout the experiment using a −10 mV hyperpolarizing step (100 ms) from −70 mV. Recordings were excluded if these values changed by more than 15% over the course of the experiment. Patch pipettes (2-4 MΩ) were filled with internal solution containing (in mM): 125 KCl, 2.8 NaCl, 2 MgCl_2_, 2 ATP-Na^+^, 0.3 GTP, 0.6 EGTA, and 10 HEPES (pH 7.25-7.28; 265-280 mOsm; junction potential at 30°C = 3.4 mV). In some experiments, guanosine 5’-O-thio-diphosphate (GDPβS; 1 mM) was substituted for GTP in the internal solution and cells were held for at least 15 minutes prior to depolarization to allow for intracellular diffusion. Evoked synaptic responses were elicited using a bipolar stainless-steel stimulating electrode. For inhibitory postsynaptic current (IPSC) recordings, the stimulating electrode was placed 200-500 µm caudal to the recording site. For excitatory postsynaptic current (EPSC) recordings, the stimulating electrode was placed <200 µm from the recorded cell within the VTA. Both IPSCs and EPSCs were evoked at 0.1 Hz using 100 µsec current pulses. To trigger somatodendritic release of CCK, dopamine neurons were depolarized in voltage-clamp to −40 mV for 6 min. For simultaneous recordings of two dopamine neurons, the cell somata were selected to be within ∼100 µm of one another, and IPSCs were evoked simultaneously in both neurons using a single stimulating electrode.

### Data analysis

IPSC and EPSC peak amplitudes were measured and normalized to baseline values by dividing each response by the mean amplitude during the 10 min baseline period. Normalized responses were binned in 2-min intervals and used to generate averaged time-course plots. The expression of LTP or LTD was determined by comparing the mean synaptic current amplitudes measured over the 5 min baseline period before depolarization or CCK application with those measured over the 5 min period after the manipulation, as indicated in each experiment.

GraphPad Prism was used for statistical analysis. Sample sizes were based on our previous experiments and related literature. Data are presented as mean ± SEM. Normality of data distributions was assessed using the Shapiro-Wilk test. All data met normality assumptions and were analyzed using parametric paired *t* tests. Statistical significance was defined as p<0.05.

## Results

### Somatodendritic release of CCK induces long term potentiation of GABAergic IPSCs

We previously demonstrated that mild depolarization of a VTA dopamine neuron for several minutes induced long-term potentiation (LTP) of electrically evoked inhibitory postsynaptic currents (IPSCs), suggesting that a synapse-modifying signaling molecule is released during depolarization. This LTP was blocked by the CCK2 receptor antagonist, LY225910, consistent with the idea that CCK release from somatodendritic regions is required for the induction of this form of plasticity (Martinez Damonte et al., 2023). However, as neuropeptides may have long half-lives in a brain slice, it remained unclear whether sustained CCK signaling is necessary for the maintenance of the potentiation, or whether transient CCK release is sufficient to induce long term synaptic plasticity.

To address this question, identified VTA dopamine neurons were voltage-clamped and evoked GABA_A_ IPSCs were recorded for at least 10 minutes. After establishing a stable baseline, the dopamine neuron was depolarized to −40 mV for 6 min to trigger CCK release. Twenty minutes after depolarization the CCK2 receptor antagonist, LY225910 (1 mM), was then bath-applied. We found that blocking CCK2 receptors after LTP induction failed to reverse the potentiation, (Figure 1A-D), as expected if transient CCK release during depolarization is sufficient to drive long term plasticity at GABAergic inputs onto VTA dopamine neurons.

**Figure 1.**
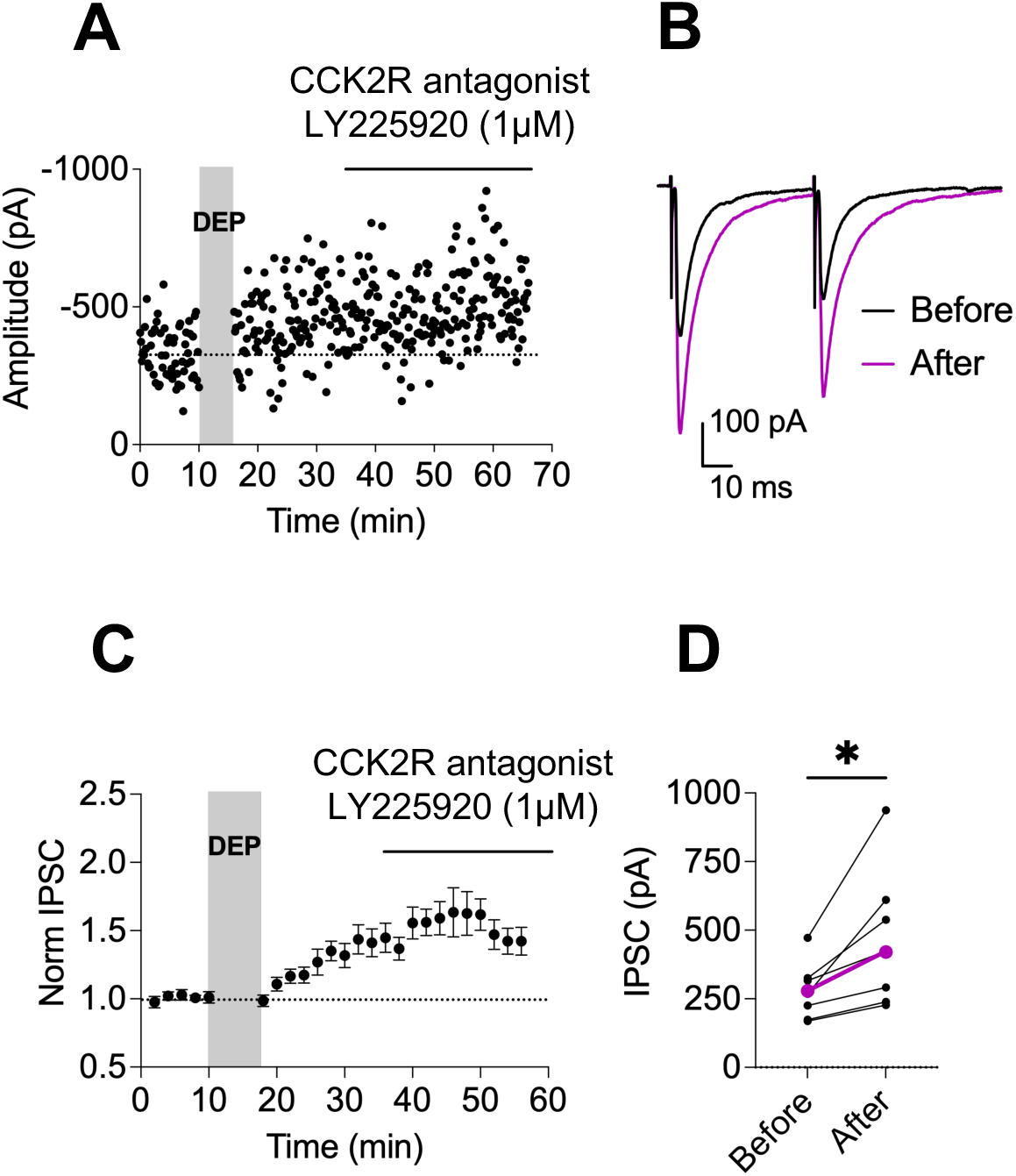
Somatodendritic release of CCK induces long-term potentiation of GABAergic IPSCs. GABAergic IPSCs were electrically evoked in voltage-clamped dopamine neurons in the VTA slice. CCK release was triggered by depolarization to −40 mV for 6 minutes (DEP, gray shaded area). The CCK2R antagonist, LY225910 was bath applied 20 minutes after dopamine neuron depolarization. (A) Representative recording showing IPSC amplitudes and LTP following depolarization, and subsequent CCK2R blockade. (B) Averaged IPSCs from this experiment during the 10 min baseline period (black, before) depolarization and 10 min after LY225910 application (purple, after)(46-56 min). (C) Time course of normalized IPSC amplitudes (n = 8 cells, 7 mice). (D) Averaged IPSC amplitudes 5 min before depolarization and during the final 5 min of recording (51-56 min; n = 8 cells, 7 mice). Paired *t* test, *p* = 0.01, *df* = 7. Colored symbols/lines represent the mean. Error bars indicate SEM.

### CCK signaling in the postsynaptic dopamine neuron is not required for LTP

We next examined whether CCK2 receptors on the postsynaptic dopamine neurons are required for LTP. Previous fluorescence in situ hybridization revealed no detectable mRNA expression of either CCK1 or CCK2 receptors in the VTA, suggesting that CCK receptor transcripts may be localized in distal presynaptic neurons rather than in local dopaminergic neurons. Here, we functionally tested whether CCK2Rs are present on the postsynaptic dopamine neurons that release CCK. CCK2 receptors are G protein-coupled receptors that mainly signal through Gαq/11 and, in some contexts, Gαi proteins (Zhang et al., 2021). We disrupted all G protein signaling in the recorded dopamine neuron by including GDPβS (1 mM) in the recording pipette (Figure 2A-E). However, even with intracellular blockade of G protein-dependent signaling, depolarization of the dopamine neuron potentiated evoked IPSCs, indicating that postsynaptic CCK2R activation is not required for the expression of this form of LTP, and that instead the CCK2Rs are most likely localized to presynaptic GABAergic terminals.

**Figure 2.**
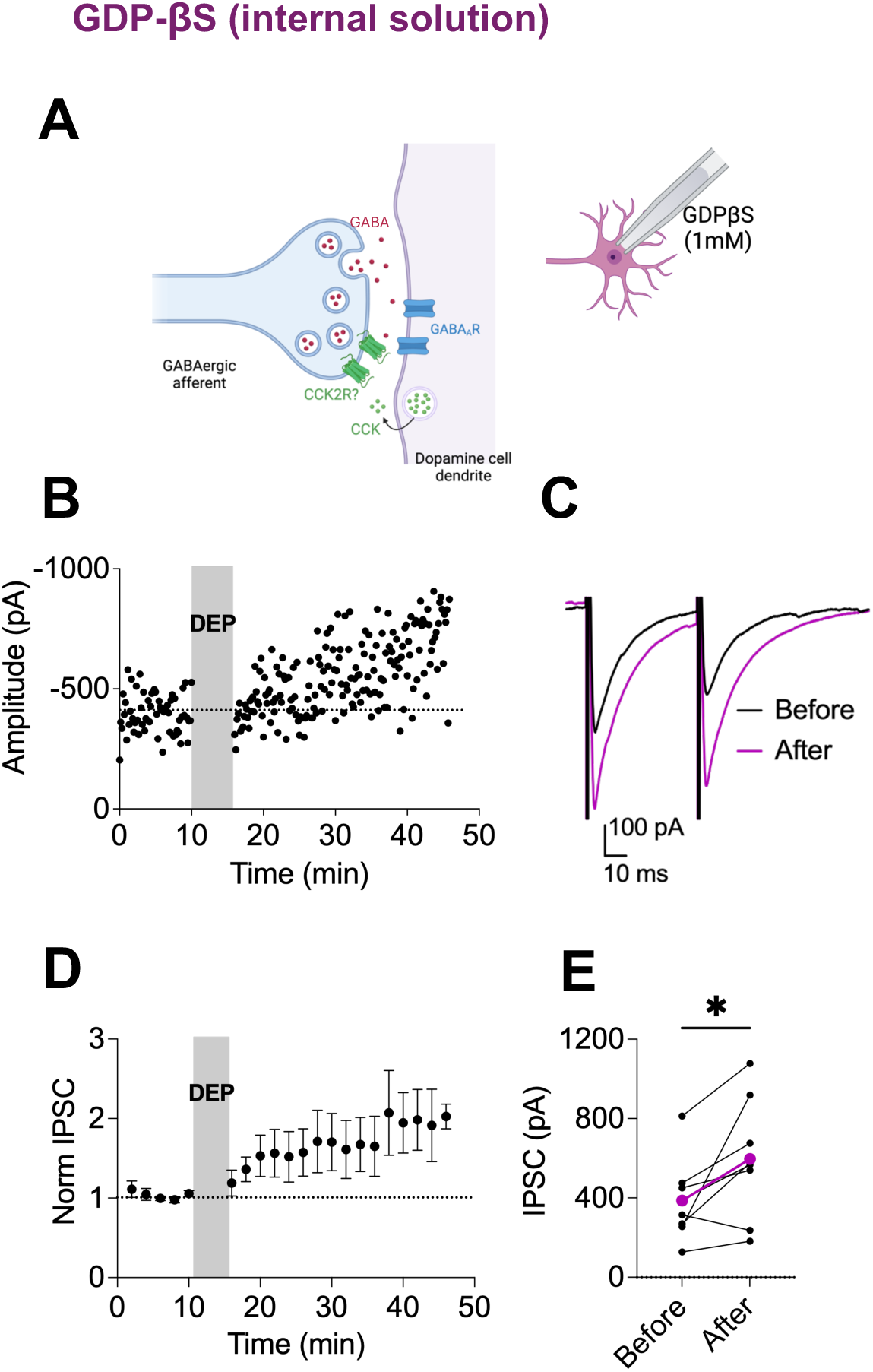
CCK receptor signaling in the postsynaptic dopamine neuron is not required for LTP. (A) Schematic illustrating somatodendritic release of CCK from a VTA dopamine neuron that induces LTP of GABAergic synaptic inputs via activation of CCK2R. GDP-bS was included in the recording pipette to block postsynaptic G-protein signaling in the dopamine neuron. (B) GABAergic IPSCs were electrically evoked in voltage-clamped dopamine neurons in the VTA slice, as in Figure 1. Representative experiment showing LTP following depolarization with 1 mM GDP-bS included in the recording pipette internal solution. (C) Average IPSCs during the 10 min baseline period (black) and 20-30 min after depolarization (purple). (D) Time course of normalized IPSC amplitudes showing LTP in cells recorded with 1 mM GDP-bS (n = 8 cells, 6 mice). (E) Average IPSC amplitudes during 5 min before and 5 min after (31-36 min) depolarization; n = 8 cells, 6 mice. Paired *t* test, *p* = 0.03, *df* = 7. Colored symbols/lines represent the mean. Error bars indicate SEM.

### Kappa opioid receptor activation on presynaptic GABAergic terminals prevents CCK-induced LTP

In the midbrain, kappa opioid receptor (KOR) activation has been reported to inhibit electrically evoked D2 receptor-mediated dopamine IPSCs, perhaps by preventing somatodendritic dopamine release (Ford et al., 2007). We therefore hypothesized that KOR activation also might prevent somatodendritic CCK release. KORs are prominent modulators of dopamine neuron activity, regulating tonic firing, synaptic strength, and influencing behavioral responses to psychostimulants such as cocaine (Margolis et al., 2005; Ford et al., 2006; Valdez et al., 2007; Bruchas and Chavkin, 2010; Ebner et al., 2010; Chavkin, 2013; Chefer et al., 2013; Ehrich et al., 2015; Whitfield et al., 2015), and KORs are highly expressed in the VTA, which receives dense dynorphinergic innervation from multiple afferent sources (Fallon et al., 1985; Poulin et al., 2009; Baimel et al., 2017; Abraham et al., 2022; Tejeda et al., 2022). To test whether KOR activation prevents somatodendritic CCK release, we bath applied the KOR agonist U69593 (200nM) throughout the experiment, reasoning that if KOR activation prevented CCK release, depolarization-induce LTP would not be induced. After a baseline recording of evoked IPSCs, dopamine neurons were depolarized to induce CCK release. Indeed, activation of KORs prevented CCK-dependent LTP of IPSCs (Figure 3A-E), indicating that KOR signaling disrupts CCK-dependent potentiation of IPSCs. We next asked whether the inhibitory effect on LTP results from a postsynaptic block of CCK release or signaling, or instead from a presynaptic action of KORs. Because KORs are G protein-coupled receptors, a mechanism dependent on postsynaptic KORs would require intact G protein signaling in dopamine neurons. However, when we included GDPβS (1 mM) in the recording pipette to block postsynaptic GPCR signaling, this did not rescue LTP in the presence of the KOR agonist (Figure 3F-I). These findings indicate that postsynaptic KORs are not those mediating the block of LTP. Instead, our data are consistent with a presynaptic action of KORs that opposes CCK2 receptor-dependent signaling at GABAergic terminals.

**Figure 3.**
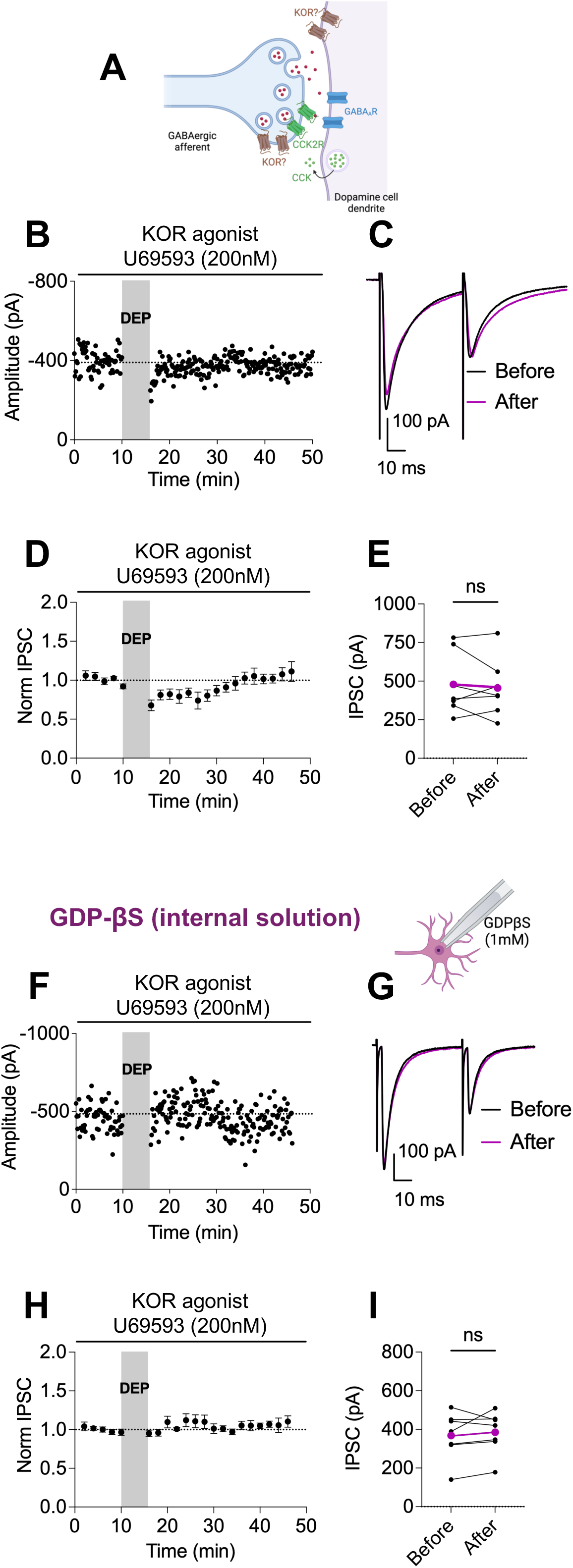
Presynaptic kappa opioid receptor activation blocks CCK-induced LTP (A) Schematic illustrating activation of KOR and their potential modulation of CCK-induced LTP at GABAergic synapses through pre- or postsynaptic mechanisms. (B) GABAergic IPSCs were electrically evoked in voltage-clamped dopamine neurons in the VTA slice, as in Figure 1. The KOR agonist U69593 was bath applied throughout the experiment. Representative time course of IPSCs, depolarization-induced LTP is blocked following KOR activation. (C) Average IPSCs during the 10 min baseline period (black) and during 20-30 min after depolarization (purple). (D) Time course of normalized IPSC amplitudes (n = 7 cells, 6 mice). (E) Average IPSC amplitudes during 5 min before and 5 min after (31-36 min) depolarization; n = 7 cells, 6 mice. Paired *t* test, *p* = 0.56, *df* = 6. Colored symbols/lines represent the mean. Error bars indicate SEM. (F-I) VTA dopamine cells were recorded with 1 mM GDP-βS included in the internal solution for at least 15 minutes before depolarization. (F) Representative time course of IPSCs in the presence of KOR agonist (U69593, 200nM). (G) Average IPSCs during the 10 min baseline period (black) and during 20-30 min after depolarization (purple). (H) Time course of normalized IPSC amplitudes (n = 7 cells, 5 mice). (I) Average IPSC amplitudes during 5 min before and 5 min after (41-46 min) depolarization; n = 7 cells, 5 mice. Paired *t* test, *p* = 0.56, *df* = 6. Colored symbols/lines represent the mean. Error bars indicate SEM.

### CCK induces long term depression of glutamatergic excitatory postsynaptic currents in dopamine neurons

Given that somatodendritic CCK release from DA neurons modulates GABAergic synapses, we next asked whether CCK also affects excitatory synaptic transmission. We first bath-applied exogenous CCK while recording pharmacologically isolated evoked EPSCs and found that CCK caused synaptic depression (Figure 4A-D). To determine whether endogenous CCK release is sufficient to drive this, we next tested whether depolarization-induced CCK release also elicits LTD of EPSCs. Consistent direct CCK application, depolarization also resulted in depression of excitatory synaptic transmission (Figure 4E-H). Together our data show that CCK exerts a dual modulatory role by potentiating inhibitory synapses while suppressing excitatory synaptic transmission onto dopamine neurons.

**Figure 4.**
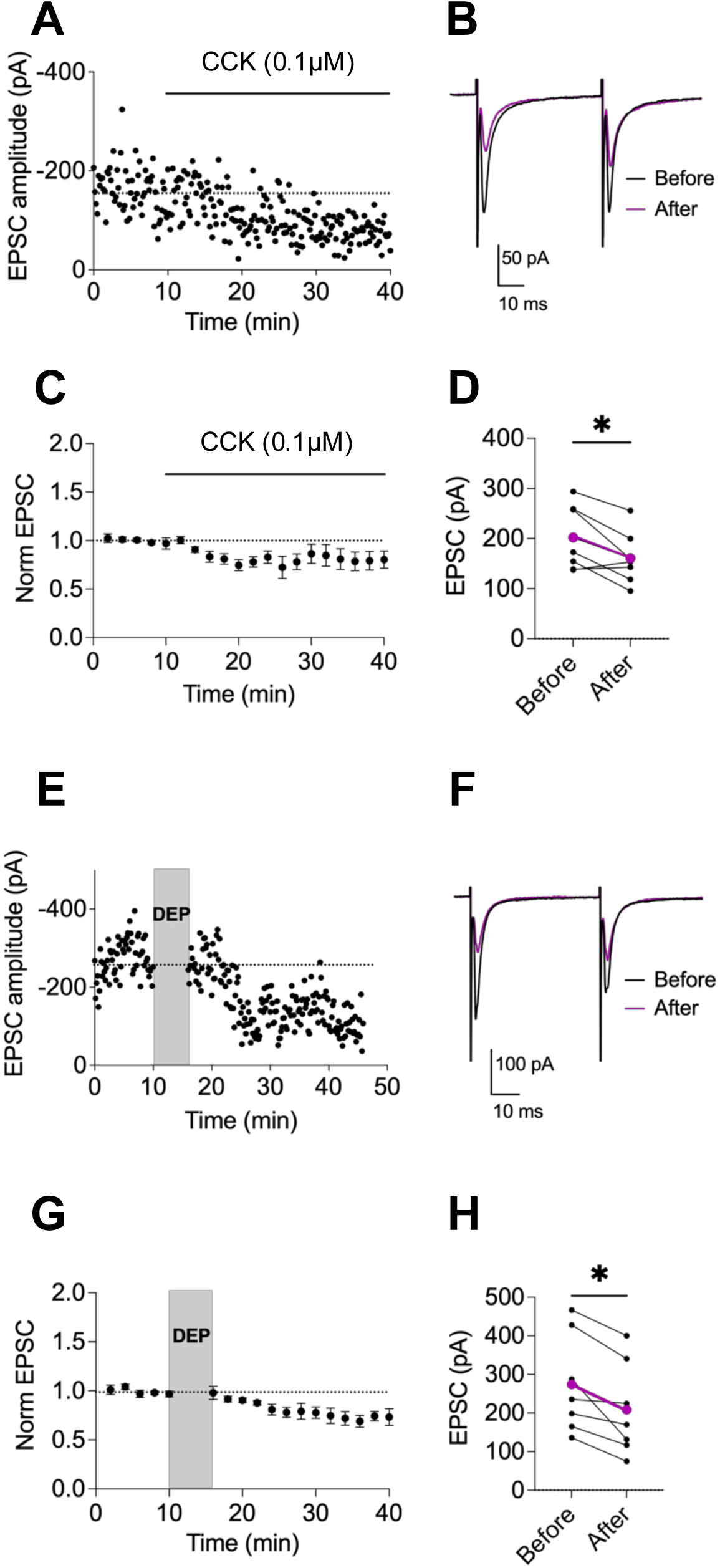
CCK induces LTD of glutamatergic EPSCs in dopamine neurons. Glutamatergic EPSCs were pharmacologically isolated and electrically evoked in voltage-clamped dopamine neurons in the VTA slice. (A-D) CCK (0.1 µM) was bath applied as noted after a stable 10 min baseline of EPSCs. (A) Representative experiment recording EPSCs showing synaptic depression following CCK application. (B) Average EPSCs from (A) during the 10 min baseline (black) and 20-30 min after CCK application (purple). (C) Time course of normalized EPSC amplitudes before and during CCK (n = 7 cells, 5 mice). (D) Average EPSC amplitudes during the 5 min before and 5 min after (25-30 min) CCK application; n = 7 cells, 5 mice. Paired *t* test, *p* = 0.03, *df* = 6. Colored symbols/lines indicate the mean. Error bars represent SEM. (E-H) Depolarization was used to release endogenous CCK; no exogenous CCK was added. (E) Representative experiment showing LTD of evoked EPSCs following depolarization (DEP, gray shaded area). (F) Average EPSCs during the 10 min baseline period (black) and 20-30 min after depolarization (purple). (G) Time course of normalized EPSC amplitudes before and after depolarization; n = 7 cells, 6 mice). (H) Average EPSC amplitudes during 5 min before and 5 min after (31-36 min) depolarization; n = 7 cells, 6 mice. Paired *t* test, *p* = 0.03, *df* = 5. Colored symbols/lines indicate the mean. Error bars represent SEM.

### CCK release from a single dopamine neuron induces LTP at GABAergic synapses in neighboring cells

Neuropeptides can diffuse from their release sites to signal through their receptors; however, the spatial range of peptide signaling in the brain remains difficult to predict due to the impact of peptidases, the extracellular matrix and the geometry of extracellular spaces (Xiong et al., 2022). Recent studies using genetically encoded fluorescence sensors have demonstrated that neuropeptides can activate receptors located more than 100 µm from release sites (Xiong et al., 2022; Dong et al., 2024). To examine the spatiotemporal dynamics of CCK signaling, we simultaneously recorded from pairs of neighboring VTA dopamine neurons less than 100 µm apart. IPSCs were then electrically evoked and recorded in both neurons (Figure 5A-B). After stable response baselines were established, only one of the two cells was depolarized to −40 mV for 6 min to induce somatodendritic CCK release. By recording the IPSCs from the non-depolarized neuron, we could assess whether CCK released in the region can affect inhibitory synapses on neighboring neurons. We found that depolarization of a single dopamine neuron was sufficient to induce LTP at synapses on the neighboring, non-depolarized dopamine neuron (Figure 5C-F). These results indicate that CCK released from one neuron spreads and can modulate inhibitory synapses on neighboring cells that have not been excited themselves.

**Figure 5.**
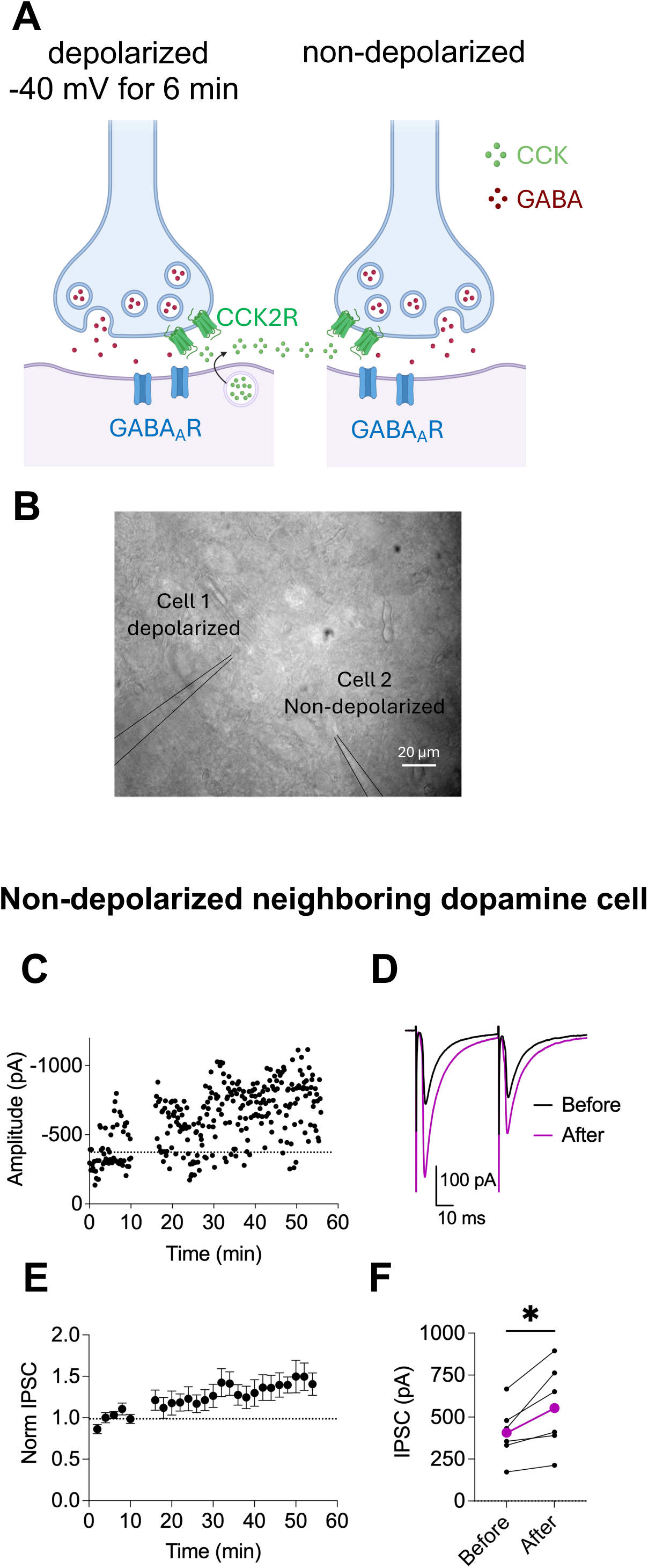
Somatodendritic CCK release from a single dopamine neuron induces LTP of GABAergic synapses in neighboring cells. (A) Schematic illustrating somatodendritic release of CCK from a depolarized VTA dopamine neuron and its diffusion to a synapse on a neighboring cell. (B) Representative image of dual whole-cell recordings from two neighboring neurons in the VTA. Cell 1 was depolarized to induce CCK release, while evoked IPSCs were recorded from the second, non-depolarized cell. (C) Representative time course of IPSC amplitudes showing LTP following depolarization of the neighboring cell at 10-16 minutes. (D) Average IPSCs from (C) during the 10 min baseline period and 10 min (46-56 min) after depolarization from neighboring cell. (E) Time course of normalized IPSC amplitudes from non-depolarized cells (n = 6 cells, 6 mice). (F) Average IPSC amplitudes from non-depolarized cells 5 min before depolarization of the neighboring cell and during the final 5 min of recording (41-46 min; n = 6 cells, 6 mice). Paired *t* test, *p* = 0.03, *df* = 5. Colored symbols/lines indicate the mean. Error bars represent SEM.

## Discussion

Here we have demonstrated that somatodendritic release of CCK from VTA dopamine neurons is sufficient to induce long-lasting and spatially extensive synaptic plasticity. Brief depolarization of a dopamine neuron triggers CCK release that in turn produces long-term potentiation of GABAergic inhibitory synapses on that neuron that persists even when CCK receptors are blocked post-depolarization. Surprisingly, LTP was elicited not only in the depolarized neuron, but also at synapses on a neighboring dopamine neuron, suggesting significant spread of the CCK within the neuropil. CCK or depolarization of a dopamine neuron also simultaneously induced LTD at glutamatergic excitatory inputs. Together, these complementary forms of plasticity are expected to suppress dopamine neuron excitability in the VTA beyond the region of the releasing neuron.

### Significant spatial spread of CCK signaling within the VTA

Our paired-recording experiments demonstrate that CCK released from a single dopamine neuron is sufficient to induce long-term synaptic plasticity in neighboring neurons with somas located at least ∼100 µm away, the greatest distance we tested. This range is notable given the compact size of the VTA, which in mice spans on the order of several hundred micrometers along its medial–lateral and dorsal–ventral axes (Beier et al., 2015). However, dopamine neurons within this region in the mouse possess extensive and highly branched dendritic arbors that can extend hundreds of micrometers and often cross subregion boundaries (Montero et al., 2021). The combination of large dendritic fields and peptide diffusion suggests that somatodendritic CCK release may influence a substantial fraction of nearby dopamine neurons, coordinating plasticity across local ensembles rather than acting in a strict cell- or synapse-autonomous manner.

### Somatodendritic CCK promotes LTD of excitatory afferents and LTP of inhibitory afferents

Long-term changes in synaptic strength at both inhibitory and excitatory synapses on VTA neurons have a key role in shaping dopamine neuron output and have been strongly implicated in reward learning, motivational salience, and addiction-related behaviors (Kauer and Malenka, 2007; Pignatelli and Bonci, 2015). Importantly, we report now that blocking G protein–dependent signaling in the dopamine neuron did not prevent CCK-induced LTP, indicating that postsynaptic CCK receptor signaling in dopamine neurons is not required for LTP. Instead, it is likely that differences in CCK receptor expression and associated downstream signaling machinery at specific GABAergic afferent terminals must determine sensitivity to CCK. Which afferent populations are most strongly influenced by somatodendritic CCK release? VTA dopamine neurons receive GABAergic input from diverse sources, including local interneurons and long-range projections from the periaqueductal gray (PAG), rostromedial tegmental nucleus (RMTg), ventral pallidum, lateral hypothalamus (LH), dorsal raphe, bed nucleus of the stria terminals (BNST) and nucleus accumbens (Beier et al., 2015; Faget et al., 2016; Beier et al., 2019; Soden et al., 2020). Our previous work demonstrated that CCK-dependent LTP of GABAergic synapses in the VTA is synapse-specific; in horizontal brain slices, CCK selectively potentiates caudally-but not rostrally-evoked IPSCs following dopamine neuron depolarization (St Laurent and Kauer, 2019; Martinez Damonte et al., 2023). Moreover, inhibitory afferents from the vlPAG, but not those from the RMTg, undergo LTP following dopamine neuron depolarization (St Laurent et al., 2020).

Along with LTP at GABAergic synapses on dopamine neurons, we found that either CCK bath application or postsynaptic depolarization elicited complimentary LTD at electrically-stimulated excitatory synapses. It was previously reported that 6 minute depolarization of a dopamine neuron by itself induces LTD at these synapses, although the mechanism was unknown (Thomas et al., 2000). Our findings identify somatodendritic CCK release as a likely mechanism capable of driving the LTD. Glutamatergic inputs to VTA dopamine neurons arise from multiple regions, including the prefrontal cortex, LH, pedunculopontine tegmental nucleus, laterodorsal tegmentum, and BNST (Sesack and Grace, 2010). Further work is necessary to define comprehensively which inhibitory and excitatory afferents may be modulated when CCK is released.

Coordinated LTD of glutamatergic synapses and LTP of GABAergic synapses should reduce dopamine neuron excitability, consistent with a role for CCK as a negative feedback signal during periods of sustained activity. Somatodendritic neuropeptide release is not restricted to discrete synaptic contacts (Van Den Pol, 2012; Ludwig et al., 2017). In this framework, synapse-specific plasticity would arise not from point-to-point connectivity but from differences in receptor expression or signaling competence among nearby terminals. This mode of action contrasts with classical LTP or LTD, which are constrained by precise synaptic connectivity, and instead suggests a role for CCK in regulating local circuit state rather than pathway-specific information transfer.

### Somatodendritic release

Somatodendritic release uses molecular mechanisms distinct from those governing classical neurotransmitter release at axon terminals, indicating that these two modes of signaling can be regulated differently. Peptides and catecholamines are packaged into dense-core vesicles that are distributed throughout both dendrites and axons (Ludwig and Leng, 2006; Park and Loh, 2008); Persoon et al., 2018), and in several well-studied systems, somatodendritic release can utilize distinct intracellular Ca²⁺ signaling mechanisms and SNARE proteins (Geppert et al., 1994; Ludwig et al., 2002; Ludwig and Leng, 2006; Schonn et al., 2008; Ford et al., 2010; Mendez et al., 2011; Van Den Pol, 2012; Jackman et al., 2016; Delignat-Lavaud et al., 2022; Lebowitz et al., 2025). Somatodendritic release of dopamine has been known for many years (Geffen et al., 1976; Korf et al., 1976), and elicits relatively rapid D2 receptor-mediated IPSCs in midbrain dopamine neurons (Rice et al., 1997; Beckstead et al., 2004; Lebowitz et al., 2025). Rapid firing or bursting releases dopamine, and the D2 IPSCs are hypothesized to provide negative feedback, hyperpolarizing dopamine neurons over hundreds of milliseconds (Ford et al., 2010; Hikima et al., 2021). In contrast, many somatodendritically released neuropeptides, like CCK, act over a longer time scale of minutes or more. For example, neurotensin release from substantia nigra dopamine neurons triggers LTD of D2 IPSCs (Tschumi et al., 2022), although the synaptic depression can be reversed by application of neurotensin antagonists (Piccart et al., 2015), indicating that it requires sustained receptor activation. Somatodendritically released vasopressin and oxytocin, which autoregulate burst firing of magnocellular neurons, each produce sustained modulation lasting tens of minutes (Ludwig et al., 2017; Brown et al., 2020). In the VTA, CCK-induced LTP persists even when CCK receptors are blocked after LTP induction, indicating that CCK initiates downstream mechanisms that maintain synaptic potentiation beyond the period of receptor activation.

### Kappa opioid receptors gate somatodendritic CCK signaling

Neuropeptide signaling in the VTA occurs within a complex neuromodulatory environment that includes the KOR (Jin et al., 2023; Mohammadkhani et al., 2024; Xu et al., 2024; Bernstein et al., 2025). KORs are highly expressed in the VTA and are activated by dynorphin afferents, where they play a well-stablished role in aversion, depression-like behaviors and stress-induced reinstatement of drug seeking (Chefer et al., 2013; Polter et al., 2014; Ehrich et al., 2015; Polter et al., 2017; Margolis and Karkhanis, 2019; Martinez Damonte et al., 2025). Here we find that activation of KORs also selectively blocks CCK-induced inhibitory LTP, likely via a presynaptic mechanism, identifying an interaction between two peptide signaling systems that modulate dopamine neuron excitability. Activation of KORs prevented depolarization-induced LTP of GABAergic inputs to dopamine neurons, even when postsynaptic G-protein signaling was blocked. Rather than acting on postsynaptic CCK receptors on dopamine neurons to suppress CCK release, our results support the idea that KORs act on GABAergic terminals to oppose CCK-dependent signaling. Previous work from our lab and others found that KORs also block a distinct form of LTP at GABAergic synapses, over a lengthy time frame of days (Graziane et al., 2013; Polter et al., 2017; Martinez Damonte et al., 2025). Moreover, KOR activation also inhibits electrically-evoked dopamine D2 receptor-mediated IPSCs, possibly by preventing somatodendritic dopamine release, removing an inhibitory feedback mechanism regulating dopamine cell firing (Ford et al., 2007). Given the prominent role of KOR signaling in stress and aversive states, our current results add to the hypothesis that activation of KORs within the VTA removes multiple natural brakes on dopamine neuron excitability, providing a potential mechanism by which dynorphin released during stressors may constrain excitability and plasticity within dopamine circuitry and reshape motivational processing.

### Somatodendritic CCK release in vivo

While early studies in rats reported that CCK transiently increases dopamine cell firing (Skirboll et al., 1981; Crawley, 1989; Tanaka et al., 1994; Hamilton and Freeman, 1995), others reported that CCK reduces dopamine cell firing both in vitro and in vivo (Brodie and Dunwiddie, 1987; Stittsworth Jr and Mueller, 1990; Martinez Damonte et al., 2023). Our previous work demonstrated that intra-VTA infusion of CCK in vivo reduces dopamine cell activity and food consumption, with the magnitude of dopamine cell inhibition correlated with the reduction in feeding behavior (Martinez Damonte et al., 2023). During feeding or other behaviors when dopamine neurons exhibit prolonged activity, this mode of peptide release may be preferentially engaged. Depending on the specific excitatory and inhibitory afferents that CCK modulates, coordinated plasticity could reduce dopamine cell excitability over a time course of minutes to hours. We speculate that somatodendritic CCK signaling may act to constrain VTA dopamine output during sustained behavioral states such as feeding.

## Acknowledgments

The authors would like to thank Joanna Stralka and Valentina Knapp for excellent technical assistance, and Dr. Christine Egjeberg, Dr. Charlotte Luff, and Valentina Knapp for suggestions on the manuscript.

